# Hydraulic conductance and the maintenance of water balance in flowers

**DOI:** 10.1101/030122

**Authors:** Adam B. Roddy, Craig R. Brodersen, Todd E. Dawson

**Affiliations:** Department of Integrative Biology, University of California, Berkeley, CA, USA; School of Forestry & Environmental Studies, Yale University, New Haven, CT, USA

**Keywords:** angiosperms, flower, hydraulic conductance, vein density, water balance

## Abstract

Flowers face desiccating conditions, yet little is known about their ability to transport water. We quantified variability in floral hydraulic conductance (*K_flower_*) for 20 species from 10 families and related it to traits hypothesized to be associated with liquid and vapor phase water transport. Basal angiosperm flowers had trait values associated with higher water and carbon costs than monocot and eudicot flowers. *K_flower_* was coordinated with water supply (vein length per area, *VLA*) and loss (minimum epidermal conductance, *g_min_*) traits among the magnoliids, but was insensitive to variation in these traits among the monocots and eudicots. Phylogenetic independent contrast (PIC) correlations revealed that few traits had undergone coordinated evolution. However, *VLA* and the desiccation time (*T_des_*), the quotient of water content and *g_min_*, had significant trait and PIC correlations. The near absence of stomata from monocot and eudicot flowers may have been critical in minimizing water loss rates among these clades. Early-divergent, basal angiosperm flowers maintain higher *K_flower_* due to traits associated with high rates water loss and water supply, while monocot and eudicot flowers employ a more conservative strategy of limiting water loss and may rely on stored water to maintain turgor and delay desiccation.

## Introduction

Flowers are often considered the hallmark of angiosperm evolution. Their appearance enabled the evolution of more intimate, specialized interactions with animal pollinators than had previously been possible (Kölreuter 1761; Sprengel 1793; Darwin 1888; Fenster et al. 2004; Crepet & Niklas 2009). Despite the influence of flowers on plant-pollinator networks (Memmott & Waser 2002), the generation and maintenance of biodiversity (Stebbins 1970; Dodd et al. 1999), and ecosystem services (Costanza et al. 1997), the fundamental relationships between floral structure and physiological function have been relatively ignored, even though non-pollinator agents of selection, such as physiological costs, often oppose the effects of pollinators (Galen et al. 1999; Galen 1999; Strauss & Whittall 2006; Lambrecht 2013).

Like leaves, flowers are terminal structures often located in the hottest, driest parts of the plant canopy, but flowers and leaves have experienced different selective forces. Leaves are a critical component in the plant hydraulic pathway and have a large influence on the terrestrial water cycle (Hetherington & Woodward 2003). Because carbon assimilation is mechanistically linked to plant transpiration (Brodribb & Feild 2000; Sack & Holbrook 2006; Brodribb et al. 2007), moving water from the roots to the leaves requires coordination in the structural traits governing water flow through each component in this pathway to prevent declines in water content that could irreversibly prohibit hydraulic function (e.g. cavitation; Drake et al. 2015; Skelton et al. 2015). Angiosperm leaves are particularly capable of efficiently transporting water because of high leaf vein densities (‘vein length per area,’ VLA) that move water closer to the sites of evaporation (Brodribb et al. 2007; Brodribb et al. 2010; Feild & Brodribb 2013; Buckley 2015), minimize changes in leaf water content (Noblin et al. 2008; Zwieniecki & Boyce 2014), and, ultimately, increase growth rates (Berendse & Scheffer 2009). The fundamental constraint of maintaining water balance by matching liquid water supply to vapor loss determines the range of possible leaf designs, from the organization of cells within leaves to the shapes and sizes of leaves themselves (Sack et al. 2008; Sack et al. 2012; John et al. 2013; Li et al. 2013).

Unlike leaves, flowers are relatively ephemeral and assimilate little carbon (but see Galen et al. 1993 for an important exception) though they still may transpire significant amounts of water (Roddy & Dawson 2012; Teixido & Valladares 2014). Instead, they promote pollen dispersal with animal- and wind-pollinated flowers differentiating along a spectrum of carbon and water investment. Because flowers often face desiccating conditions that would lead to wilting and prevent successful pollination, they must maintain water balance and turgor throughout anthesis to attract pollinators. At least part of this hydraulic pathway may need to remain functional after pollination to facilitate proper fruit and seed development. Understanding how flowers do this is fundamentally important to understanding non-pollinator agents of selection and, by extension, floral evolution.

Even basic information about flower water relations, such as the mechanisms used to deliver water, are not clear. Basal angiosperm flowers from the ANA grade (*Amborella*, Nymphaeales, Austrobaileyales) and magnoliids seem to rely predominantly on continuous delivery of water by the xylem (Feild et al. 2009a, b). Such a strategy could be costly because it would require a vascular system capable of meeting potentially large transpiration demands and risky because it would require flower water potential to decline diurnally with stem water potential. In contrast, some eudicot flowers and petals tend to have higher, less negative water potentials than subtending bracts and leaves (Trolinder et al. 1993; Chapotin et al. 2003). These ‘reverse’ water potential gradients imply that for these flowers to remain hydrated, water must be imported against an apoplastic water potential gradient in the xylem. These authors have concluded that flowers are hydrated predominantly by the phloem, which is primarily responsible for the transport of photosynthates throughout the plant. In contrast to the xylem, the phloem has much higher hydraulic resistance and lower water flux rates because phloem transport occurs symplastically (Münch 1930; Nobel 1983; Windt et al. 2009; Savage et al., *in press*). However, implicating the phloem as the sole source of floral water is not necessary to explain the results from these studies; flowers can be xylem-hydrated and still maintain higher water potentials than leaves so long as they are more negative than the stem xylem (e.g. Feild et al. 2009), or they could rely on large amounts of water imported early in development (e.g. during bud expansion) and stored locally (i.e. hydraulic capacitance; Chapotin et al. 2003) that could be depleted throughout anthesis. Regardless of whether flowers have high hydraulic capacitance or rely on water delivered by the phloem, maintaining a higher water status in flowers could result in water being drawn back into the stem during the day, as has been shown to occur in fleshy fruits (Higuchi & Sakuratani 2006). The supposed dichotomy between xylem-hydration and phloem-hydration in flowers is likely not a dichotomy after all but rather a spectrum between more or less contribution from the phloem. In fruits, there has been a similar debate about the contribution of the phloem (Ho et al. 1987; Lang 1990; Greenspan et al. 1994), and the most recent evidence suggests that both the xylem and the phloem supply water, but the relative contributions of the two pathways may vary during development (Choat et al. 2009; Windt et al. 2009; Clearwater et al. 2012; Clearwater et al. 2013).

Regardless of the pathways of water entry into flowers, prior evidence suggests that there may be substantial variation in the hydraulic structure-function relationships of flowers and that much of this variation may be explained by phylogenetic history. In the present study, we quantified whole flower hydraulic conductance (*K_flower_*) for 20 species from 10 families across the angiosperm phylogeny as a first attempt at characterizing interspecific variation in hydraulic capacity. *K_flower_* has previously been measured on only one species, *Magnolia grandiflora* (Feild et al. 2009). Using leaf structure-function relationships as a starting point, we hypothesized that *K_flower_* would be mechanistically linked to xylem traits that may control water supply, such as *VLA* and the Huber ratio, and also to stomatal and epidermal traits of tepals and petals that may regulate water loss rates, such as stomatal density and size and epidermal conductance to water vapor. Positive correlations between *K_flower_* and other hydraulic traits would suggest that these traits are important in determining the efficiency of water flow through flowers. We further predicted that positive correlations between *K_flower_* and hydraulic traits, particularly xylem traits, should exist for magnoliid flowers because flowers of two of these genera, *Magnolia* and *Calycanthus*, have been shown to maintain a hydraulic connection to the stem xylem throughout anthesis (Feild et al. 2009b; Roddy et al. in prep.). If flowers from the eudicots, and possibly also the monocots, rely more heavily on a different source of water during anthesis (e.g. either delivery by the phloem or depletion of stored water), then they should not exhibit positive correlations between xylem traits and *K_flower_*. Given that much of the variation in traits may be due to differences among clades, we used correlations of phylogenetic independent contrasts (PICs) to test whether pairs of traits coevolved (Felsenstein 1985). Coevolution of traits would further imply a functional link.

## Materials and Methods

### Plant material

We collected flowering shoots from around the University of California, Berkeley, campus and from the University of California Botanic Garden during the springs of 2013 and 2014, and from the Marsh Botanical Garden, New Haven, CT, in the spring of 2015. All plants had been kept well-watered. We chose a phylogenetically diverse set of species that varied by almost two orders of magnitude in floral display size (Table 1). These species also varied morphologically, from flowers with undifferentiated perianths to those with a fully differentiated calyx and corolla and from those with free petals to those with sympetalous connation. Additionally, we included inflorescences of *Cornus florida* (Cornaceae), which have small, inconspicuous flowers but large, white bracts as their showy organs. Although the showy floral structures of this set of species are not homologous, they have a convergent function of attracting pollinators. For each species, we measured *K_flower_* on at least three flowers, and most species had low variance among individual flowers. Sample sizes for each species are shown in Table 1.

**Table 1.**
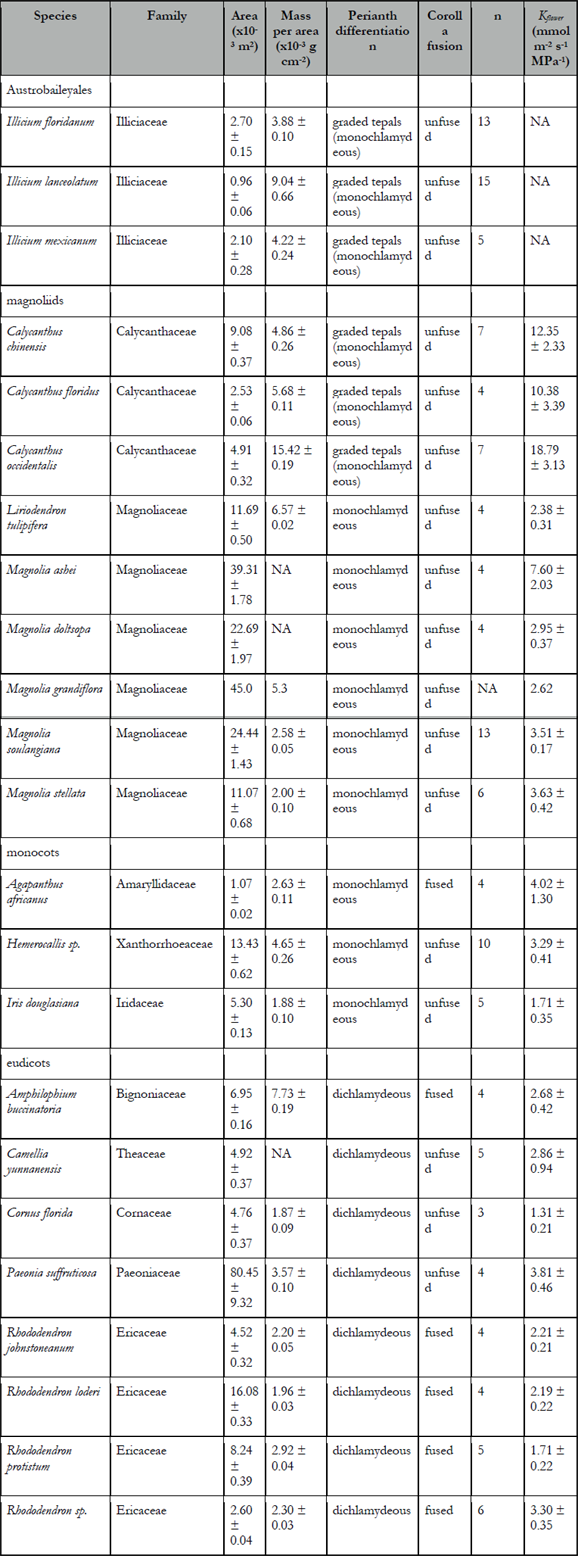
List of species sampled and their morphological traits and sample sizes (*n*) for measurements of *K_flower_*.

### Measurements of hydraulic conductance

We measured hydraulic conductance of whole, recently opened flowers using a low pressure flow meter (LPFM) under low-light, laboratory conditions (Kolb et al. 1996). We chose this method rather than the evaporative flux method because the evaporative flux method depends on maximizing boundary layer conductance. Because of the morphological complexities of flowers (i.e. unlike leaves, flowers are rarely planar), we were not confident we could maximize the boundary layer conductance to obtain realistic maximum values of *K_flower_*. However, the LPFM has the potential to clear any xylem occlusion, which we tested on a subset of species by (1) comparing flow rates when increasing the vacuum pressure to flow rates measured when decreasing the vacuum pressure and (2) by repeatedly measuring the same flower. We found no differences between flow rates measured while increasing the vacuum or while decreasing the vacuum and no significant increase in *K_flower_* with subsequent measurements (data not shown).

Flowering shoots were excised early in the morning (before 9:00 am) when stem water potentials of plants growing in this area are generally higher than -0.25 MPa. Cut shoots were immediately recut under distilled water at least one node apical to the first cut and transported back to the lab in a covered bucket to minimize water loss. Shoots were kept in water in the covered bucket for at least one hour during transport and after returning to the lab before any flowers were excised, allowing for relaxation of xylem tension. We only measured the most recently opened flowers on each plant, based on each flower’s development relative to other flowers on the shoot. Once in the lab, individual flowers were excised underwater at the pedicel base and connected to hard-walled tubing. Pedicels were glued into short lengths of tubing of various diameters using cyanoacrylate glue, and this tubing was, in turn, connected with a compression fitting to hard-walled tubing that led back to an electronic balance with resolution to 0.1 mg (Sartorius CPA225 or Practum 224-1S, Sartorius, Goettingen, Germany), on which sat a vial of dilute electrolyte solution (10 mM KCl, filtered to 0.2 um and degassed the morning of measurements). Flowers were placed in a cylindrical acrylic chamber that was attached to a vacuum pump and lined with wet paper towels. Flow rates of KCl solution into the flower from the balance were measured every 10-60 seconds depending on the absolute flow rate under 5-6 different pressures ranging from 15 to 60 kPa below ambient. At each pressure, flow rates were allowed to stabilize for 3-20 minutes and until the coefficient of variation of the last ten readings was less than 5% and the instantaneous measurements converged on the average of the last ten measurements. In practice, low absolute flow rates meant that stable averages could be reached but the coefficient of variation often remained above 5%. To determine *K_flower_*, we linearly regressed the flow rates versus pressure and removed, at most, one outlying point from the regression (Supplementary Figure 1). Flowers of most species did not have any points removed, and regressions used to calculate *K_flower_* had *R*^2^ values above 0.85. Points were removed if the relationship between pressure and flow rate was non-linear. Often, the point at the highest pressure was responsible for causing non-linearity, and this point was removed, which brought *R*^2^ values above 0.85. Immediately after measurements, we scanned the flowers to determine the projected surface area of all perianth parts, which we used to normalize hydraulic conductance to calculate *K_flower_* in units of mmol s^-1^ m^-2^ MPa^-1^. Only two species, *Amphilophium buccinatoria* and *Paeonia suffruticosa*, had a persistent, green calyx, which comprised 7.6% and 5.5%, respectively, of the total evaporative surface area. For comparison, values of *K_flower_* for *Magnolia grandiflora* reported by Feild et al. (2009b) using a different method were approximately equivalent to values produced using our method for congeneric species, and maximum measurements of *K_flower_* in the field based on transpiration rate and the water potential gradient between stem and flower for *Calycanthus occidentalis*were equivalent to those presented here using the low pressure flow meter (data not shown). We excluded *K_flower_* data for *Illicium* flowers because flowers of these species had open paths for water transport that, we believe, produced erroneously high values of *K_flower_*. Almost immediately after applying the vacuum, water seeped out of the center of the flowers or formed rapidly growing droplets on the tepal surfaces. This behavior was unique to *Illicium* flowers and justified excluding them from subsequent analyses involving *K_flower_*.

### Trait measurements

We measured two xylem traits predicted to be associated with water supply, the Huber ratio and *VLA*. The Huber ratio is the ratio of the xylem cross-sectional area in the pedicel to the evaporative surface area. Immediately after measuring *K_flower_* pedicels were free hand sectioned underwater. The sections were placed in distilled H_2_O, while floral structures (tepals, petals, sepals) were individually removed and scanned on a flatbed scanner. The pedicel cross-sections were quickly stained with phloroglucinol and imaged at 5-40x under a compound microscope outfitted with a digital camera. We measured the xylem cross-sectional area and the surface area of flowers using ImageJ (version 1.44o; Rasband 2012). Sampling for vein density was identical to Roddy et al. (2013) and briefly summarized here. To account for the high variability in vein density within a petal, we excised multiple 1-cm^2^ pieces from petals of at least four flowers. These sections were placed in 2% NaOH for clearing. Sections were rinsed briefly in distilled H_2_O and then placed in 95% ethanol. Once in ethanol, samples were briefly stained with Safranin O and imaged at 5-20x magnification under a compound microscope outfitted with a digital camera. One or two images per section from each of five to twelve sections per species were captured, and vein densities were measured using Image J (version 1.44o; Rasband 2012).

We also measured traits predicted to influence rates of water loss. These included stomatal traits (stomatal density, guard cell length length, and stomatal pore area index [SPI]) and the minimum epidermal conductance to water vapor, *g_min_*. The minimum epidermal conductance is the area-normalized conductance to water vapor, measured in the dark after stomata have been allowed to close (Kerstiens 1996). We measured *g_min_* on individual floral structures by sealing the cut edges with a thick layer of petroleum jelly and placing them in a dark box into which was placed a fan and a temperature and relative humidity sensor. For connate flowers, we measured the entire tubular structure and sealed the cut base with petroleum jelly. Structures sat on a mesh screen while the fan pulled air across the flowers inside the container. Every 5 to 20 minutes, the container was briefly opened and the structure weighed on a balance with a resolution of 0.1 mg. A regression of the linear part of this resulting curve was used to calculate *g_min_*. After approximately 10 measurements, each structure was scanned to measure its area and then placed in a drying oven for later dry mass measurement. Using the temperature, humidity, mass, and area measurements, we calculated *g_min_* and the desiccation time (*T_des_*), which we define as the time required for the structure to fully desiccate and is calculated as:

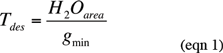

where *H_2_O_area_* is the fresh water content per area. The fresh water content was defined as the difference between the saturated mass measured immediately after excision during measurements of *g_min_* and the final dry mass, divided by the projected surface area. *T_des_* therefore has units of time. A conservative strategy of limiting water loss would be associated with a long *T_des_*. In statistical analyses, *T_des_* was log-transformed to improve normality. We also calculated the floral dry mass per area (*FMA*) from the area and dry mass measurements needed for measuring *g_min_*.

We used two methods to measure stomatal traits. First, we cleared tepal and petal sections (and, in the case of *Cornus florida*, sections of showy bracts) in 2% NaOH, rinsed them briefly in distilled H_2_O, and transferred them into 95% EtOH. Images of the epidermis were made using a compound microscope at 5-40x. We imaged 5-20 fields of view to determine stomatal densities, depending on the abundance of stomata. Second, we also made stomatal impressions using dental putty (Coltene Whaledent President Light Body). In our experience nail varnish applied directly to petals is difficult to remove while maintaining a good impression. Instead, we made nail varnish impressions of the hardened dental putty negatives and imaged the nail varnish impressions with a compound microscope. Guard cell length was determined by measuring the maximum length of at least 10 guard cells for each species with stomata. All of these measurements were made on abaxial and adaxial surfaces and averaged to calculate stomatal density on a per unit area basis. The stomatal pore area index (*SPI*) was calculated as the product of stomatal density and the square of average guard cell length, according to Sack et al. (2003).

We lacked trait data for some species but, because of the paucity of these measurements on flowers, we have chosen to include these species in the present analyses when possible. Data for *Magnolia grandiflora* were taken from Feild (Feild et al. 2009) and so lacked many of the traits we measured. Because of the limited number of flowers available for *Magnolia doltsopa*, only *K_flower_* was measured on this species, which appears exclusively in Table 1 and Figure 1.

**Figure 1.**
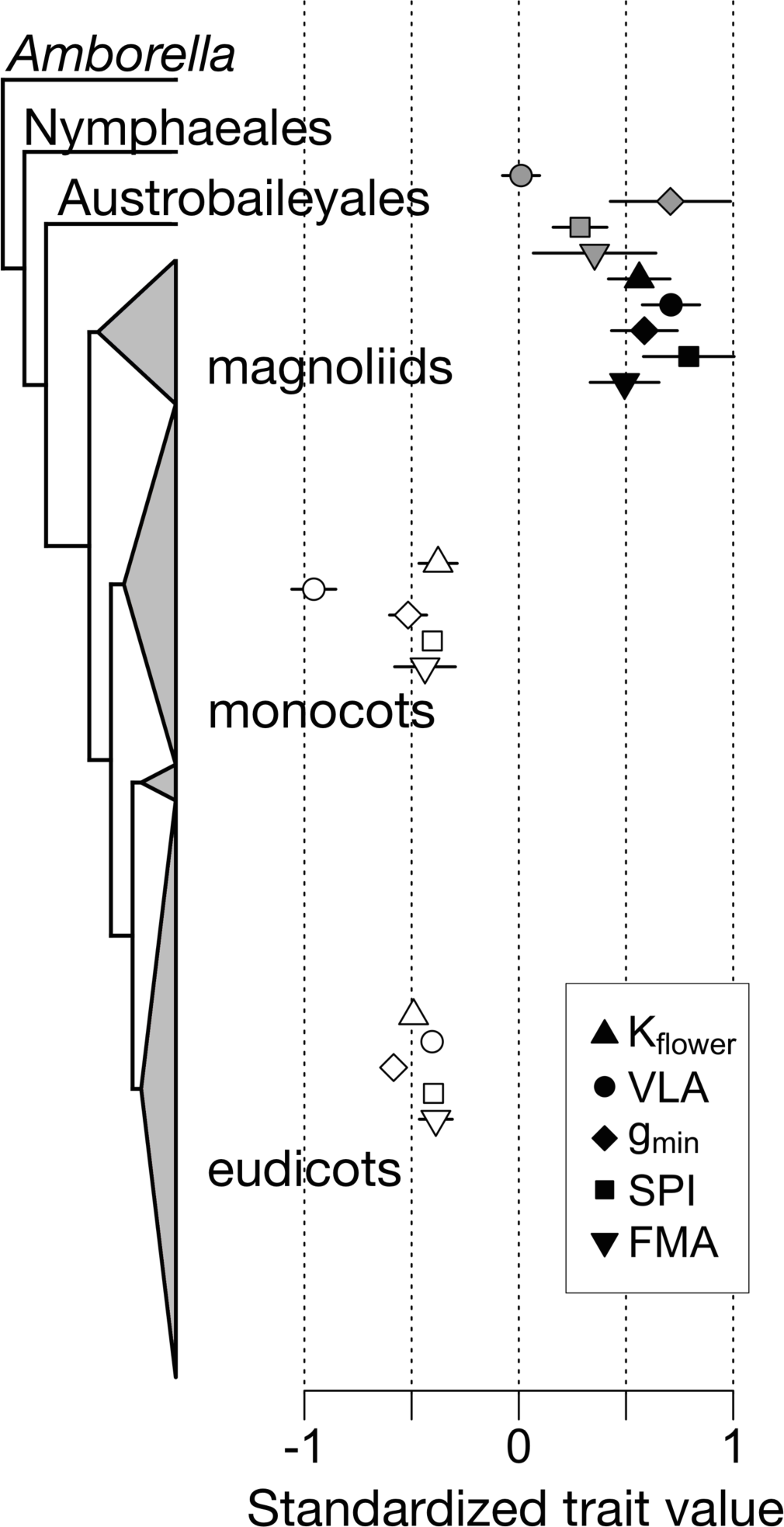
Hydraulic and structural trait variation among major angiosperm lineages. Data were scaled and normalized by their mean so that all traits are on the same scale. Points represent mean and standard error for each lineage based on the species listed in Table 1. Positive standardized trait values signify high trait values and negative standardized trait values signify low trait values.

### Statistical and phylogenetic analyses

All analyses were performed in R (v. 3.1.1; R Core Team 2012). For correlations between traits, we used the conservative non-parametric Spearman rank correlation and report the correlation coefficient (*r_s_*). Additionally, for certain relationships we fit linear regressions when appropriate and report *R^2^* values only for these linear fits. To test for correlated trait evolution, we generated a phylogeny for the species in our dataset using Phylomatic (v. 3; Webb & Donoghue 2005) with branch lengths proportional to diversity, based on the method of Grafen (Grafen 1989). Phylogenetic independent contrasts (PICs; Felsenstein 1985) were calculated using the ‘pic’ function in the R package *picante (Kembel et al. 2010).*

Because much of the variation in traits was due to large trait variation among the magnoliids, particularly the genus *Calycanthus*, we calculated trait and PIC correlations on different subsets of species. Many basal angiosperms share common ecophysiological traits (Feild et al. 2009a), and so we examined correlations only among these species and separately among only the monocots and eudicots. Similarly, we examined trait correlations among the entire dataset with and without the genus *Calycanthus* to determine the extent to which significant correlations were due to this one genus.

## Results

There was large variation among species in all traits, due mainly to differences among major clades. Basal angiosperm flowers, which include the magnoliids and *Illicium*, tended to have traits associated with higher costs in terms of both water and carbon than monocot and eudicot flowers (Figure 1). There was little variation in trait values among species in the monocots and eudicots, suggesting that these species share a physiological strategy of having low area-normalized trait values despite varying almost 75-fold in flower size in our dataset (from *Agapanthus africanus* at 10.76 cm^2^ to *Paeonia suffruticosa* at 804.53 cm^2^).

*K_flower_* varied widely among the species we studied from a mean of 1.30 mmol s^-1^ m^-2^ MPa^-1^ for *Cornus florida* inflorescences to 18.79 mmol s^-1^ m^-2^ MPa^-1^ for *Calycanthus occidentalis* flowers (Figure 2a). *C. florida* inflorescences had the lowest *K_flower_* of any species measured, despite its showy organs being bracts. Interestingly, the three magnoliid genera spanned most of the variation in *K_flower_* of all species measured, ranging from 2.38 mmol s^-1^ m^-2^ MPa^-1^ for *Liriodendron tulipifera* to 18.79 mmol s^-1^ m^-2^ MPa^-1^ for *Calycanthus occidentalis*. The monocots varied from 1.71 mmol s^-1^ m^-2^ MPa^-1^ for *Iris douglasiana* to 4.03 mmol s^-1^ m^-2^ MPa^-1^ for *Agapanthus africanus*, while the eudicots ranged from 1.30 mmol s^-1^ m^-2^ MPa^-1^ for *Cornus florida* inflorescences to 3.81 mmol s^-1^ m^-2^ MPa^-1^ for *Paeonia suffruticosa*.

**Figure 2.**
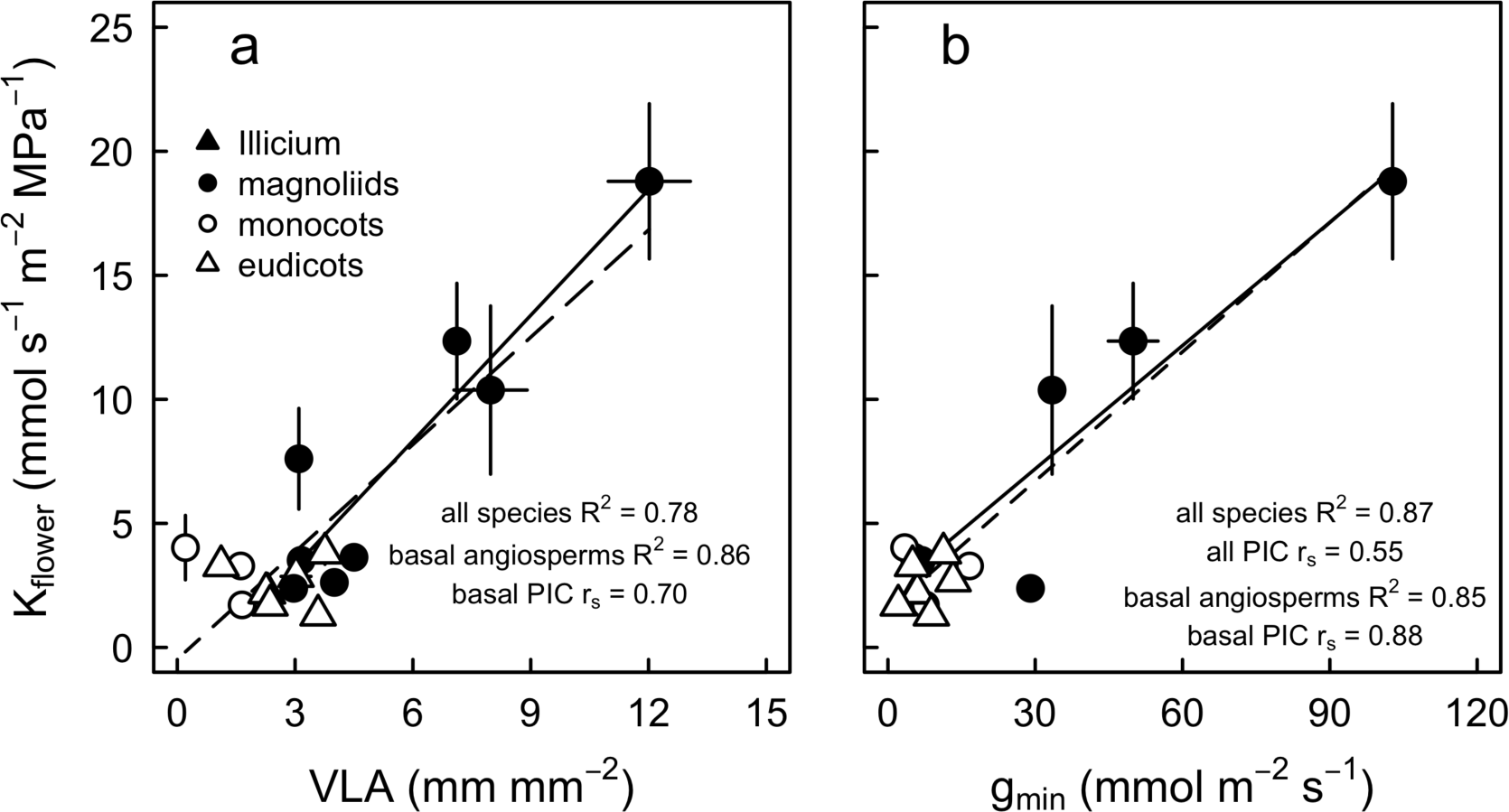
Coordination between *K_flower_* and (a) vein length per area and (b) *g_min_*. Solid lines represent linear fits only through the basal angiosperm and magnoliid points, and dashed lines represent linear fits through all species.

Other traits similarly varied approximately two orders of magnitude (Figures 1, 2). *VLA* exhibited a similar range to that seen by Roddy et al. (2013), ranging from 0.21 mm mm^-2^ for *Agapanthus africanus* to 12.02 for *Calycanthus occidentalis*. The Huber ratio ranged from 3.63 x 10^-6^ for *Cornus florida* to 2.69 x 10^-5^ for *Illicium mexicanum*, while *g_min_* values ranged from 2.10 mmol m^-2^ s^-1^ for *Rhododendron protistum* to 102.78 mmol m^-2^ s^-1^ for *Calycanthus occidentalis* tepals. Stomatal traits were similarly variable, with many species lacking floral stomata entirely, as has been reported previously (Lipayeva 1989). Among the monocots only *Iris douglasiana* had stomata (2.21 mm^-2^ and 13.08 μm in length), and among the eudicots only *Rhododendron johnstoneanum* had stomata (0.55 mm^-2^ and 21.13 μm in length). Interestingly, we did not find stomata on flowers of the other *Rhododendron* species. However, the basal angiosperms had more abundant stomata on their tepals. *Illicium lanceolatum* had the least abundant stomata (4.87 mm^-2^ and 26.63 μm in length), while *Calycanthus occidentalis* had the most abundant stomata (14.23 mm^-2^ and 16.70 μm in length).

There were surprisingly few significant correlations among traits in our dataset (Table 2). We were particularly interested in which traits were coordinated with *K_flower_*. Interestingly, only xylem traits, *VLA* and Huber ratio, were significantly correlated with *K_flower_*, although PIC correlations between *K_flower_* and *g_min_* were also significant. However, with the exception of the correlation with the Huber ratio, *K_flower_* was not correlated with anything once *Calycanthus* was excluded. *VLA* correlated with traits associated with water loss: *g_min_*, stomatal density, *SPI*, and *T_des_* (Figure 3, Table 2). However, without *Calycanthus VLA* did not correlate with any other trait, but PIC correlations between *VLA* and *T_des_* and between *VLA* and *K_flower_* were significant even without *Calycanthus*.

**Figure 3.**
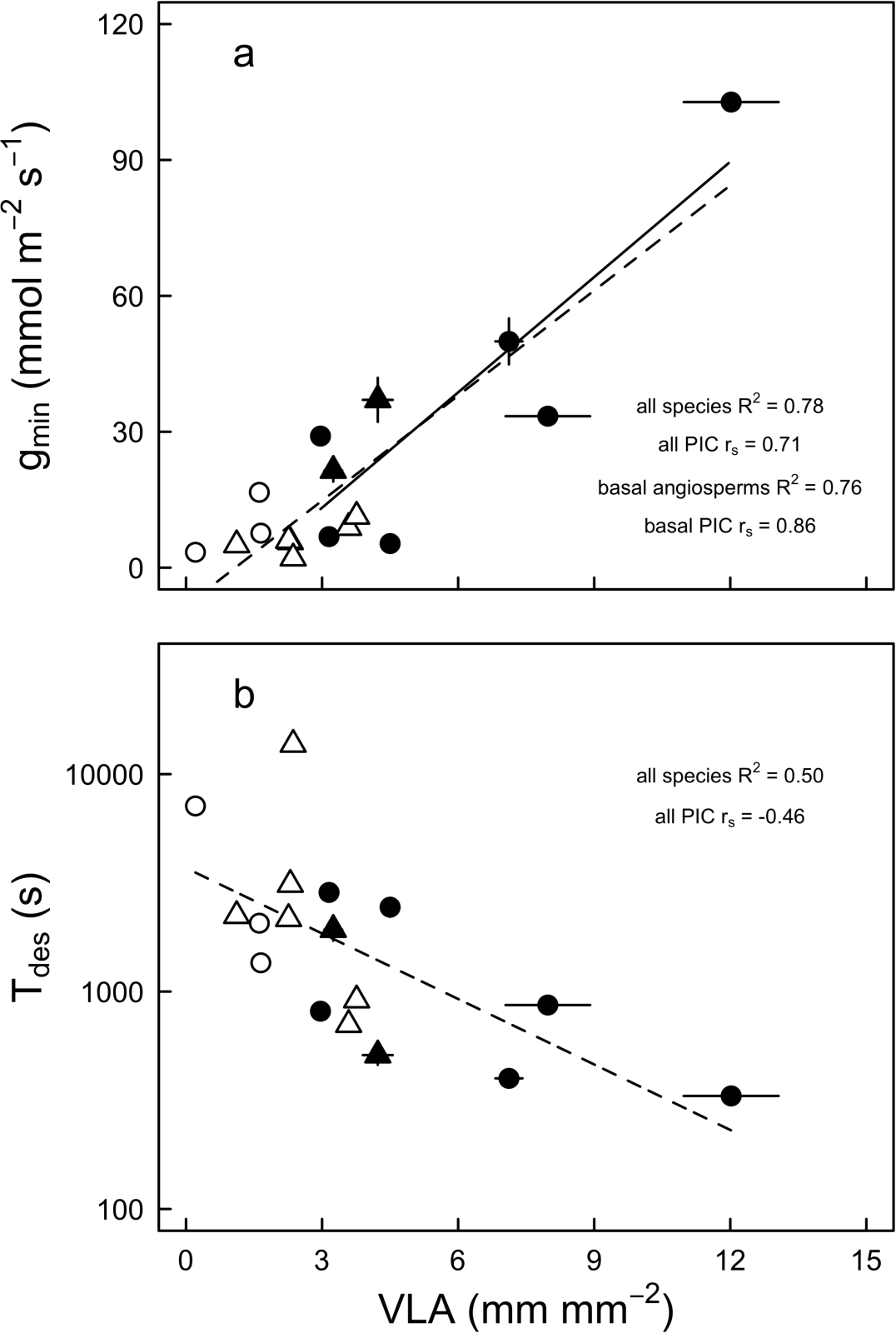
(a) Coordination between vein length per area and *g_min_*. (b) Tradeoff between vein length per area and *T_des_*. Solid lines represent linear fits only through the basal angiosperm and magnoliid points, and dashed lines represent linear fits through all species. Point symbols are the same as in Figure 2.

**Table 2.**
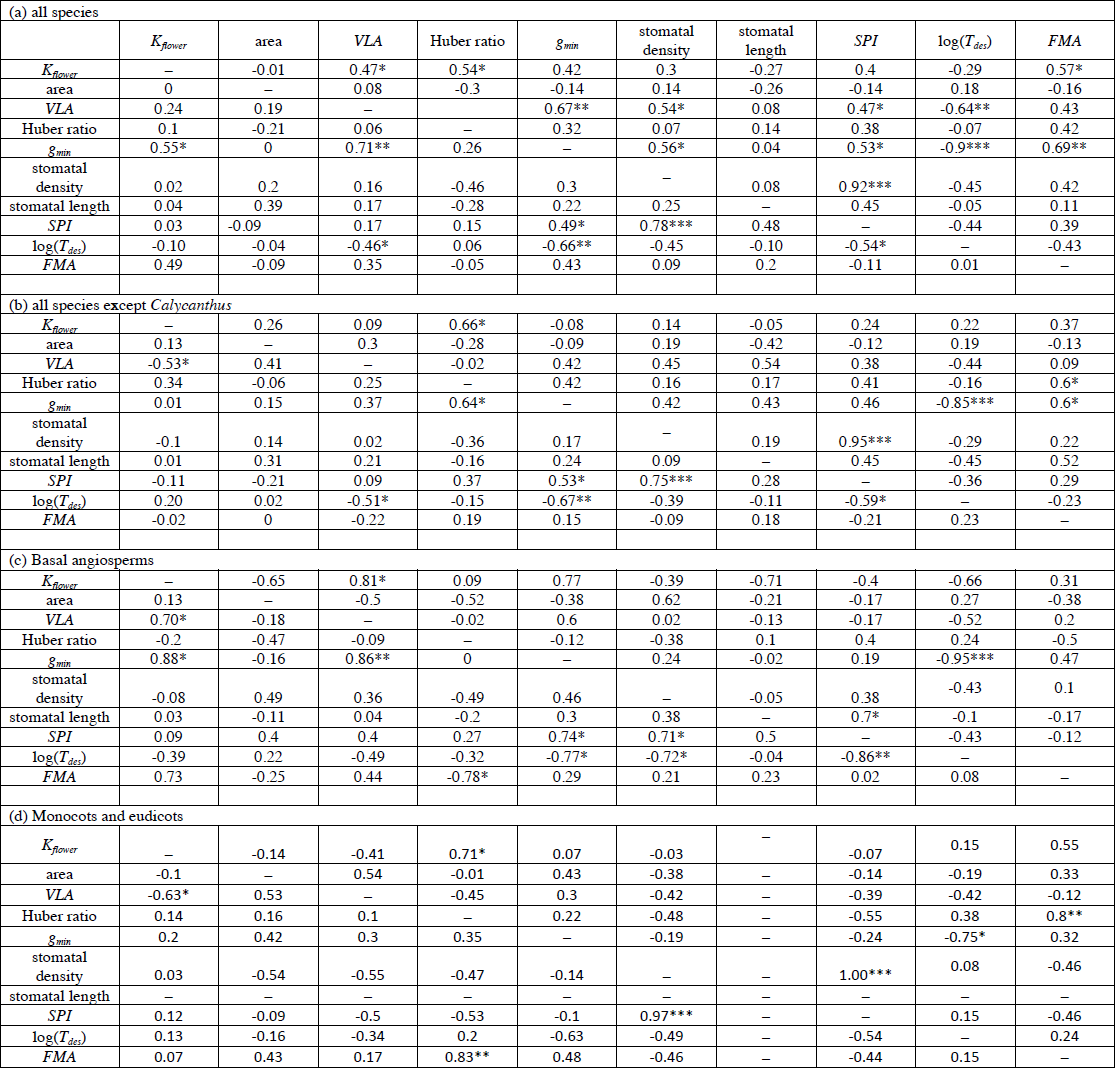
Trait and phylogenetic independent contrast correlations (a) among all species, (b) among all species except *Calycanthus*, (c) only among basal angiosperms, and (d) only among monocots and eudicots. Stomatal length is omitted from (d) because so few species had stomata. Trait correlations are in the upper triangle and phylogenetic independent contrast correlations are in the lower triangle. Spearman rank correlation coefficients are shown. Asterisks indicate significance level: *P < 0.05; **P < 0.01; ***P < 0.001.

Because most of the variation in our dataset was driven by the basal angiosperms (particularly the genus *Calycanthus*), which share many other ecophysiological traits (Feild et al. 2009), we examined trait correlations and coevolution just among these lineages. The relationship between *K_flower_* and *VLA* was stronger just among the basal angiosperms (*R^2^* = 0.86, df = 6, P < 0.001) than among all species (*R^2^* = 0.78, df = 16, P < 0.001; Figure 2), and the PIC correlation among the basal angiosperms was also significant (*r_s_* = 0.70, df = 6, P = 0.04). The correlation between *K_flower_* and *g_min_* was stronger among all species (*R^2^* = 0.87, df = 14, P < 0.001) than among the basal angiosperms (*R^2^* = 0.85, df = 4, P < 0.01), and the PIC correlations among all species (*r_s_* = 0.55, df = 13, P = 0.03) and among basal angiosperms was significant (*r_s_* = 0.88, df = 6, P = 0.02). Similarly, among the basal angiosperms, *g_min_* and *VLA* were correlated both for traits (*R^2^* = 0.76, df = 6, P < 0.01) and independent contrasts (*r_s_* = 0.86, df = 6, P < 0.01). However, while *VLA* and *T_des_* were correlated among all species both for traits (*R^2^* = 0.50, df = 15, P < 0.01; Figure 3) and contrasts (*r_s_* = -0.46, df = 14, P = 0.05), neither the trait correlation (P = 0.07) nor the contrast correlation (P = 0.20) was significant among the basal angiosperms. Although excluding *Calycanthus* from the entire dataset caused the correlation between *T_des_* and *VLA* to be insignificant, the independent contrast correlations between *T_des_* and *VLA* were significant whether or not *Calycanthus* was included for the entire dataset (Table 2).

## Discussion

Aerial plant structures often face hot, dry conditions that can lead to desiccation and impair physiological function. Despite the constraint of maintaining positive water balance in distal structures, there is substantial variation in how different structures and species avoid desiccation. For example, leaf hydraulic architecture has been optimized to efficiently transport large fluxes of water for sustained periods of time, although there is substantial variation among species, depending, for example, on leaf lifespan and dry mass investment (Simonin et al. 2012). By coordinating rates of water loss with rates of water supply, most leaves are able to avoid large declines in water content that could restrict carbon assimilation and cause irreversible hydraulic failure. Flowers, by contrast, need not assimilate carbon, so maintaining a high capacity for transporting water may not be advantageous. Furthermore, prior evidence has suggested that there may be large differences among major angiosperm clades in how flowers maintain water balance (Trolinder et al. 1993; Feild et al. 2009; Feild et al. 2009; Chapotin et al. 2003).

Almost all of the hydraulic and structural traits measured in the current study showed significant variation among species with basal angiosperms (*Illicium* and magnoliids) having trait values consistent with higher carbon and water costs (Figure 1). Basal angiosperm flowers had more abundant stomata, higher *g_min_*, higher *VLA*, higher *K_flower_*, and also higher *FMA* than flowers of the monocots and eudicots. Basal angiosperm flowers also showed strong coordination between *K_flower_*, *VLA*, and *g_min_*, suggesting that these traits may be critical in determining floral hydraulic capacity among these lineages (Figures 2, 3). Monocot and eudicot flowers, by contrast, had lower values of all traits measured except the Huber ratio, and trait values were much less variable among the monocots and eudicots. Monocot and eudicot flowers may have a common physiological trait syndrome despite large variation in other floral characters such as size, shape, and developmental origin. For example, in our dataset, monocot and eudicot flowers varied nearly 75-fold in flower size (Table 1), yet exhibited little variation in area-normalized trait values (Figures 1-3). Indeed, lower physiological and structural costs have likely relaxed the strength of non-pollinator selection on monocot and eudicot flowers and allowed morphological traits important to pollination to vary more widely.

The evolutionary pattern of trait variation (Figure 1) and prior measurements of water potentials and gas exchange (Trolinder et al. 1993; Feild et al. 2009; Feild et al. 2009; Chapotin et al. 2003) suggest that there may be fundamental differences among clades in how flowers are built and hydrated. Magnoliid flowers, including both Magnolia (Feild et al. 2009b) and *Calycanthus* (Roddy et al., in prep.), have high transpiration rates and maintain a functional connection to the stem xylem for water supply. Magnoliid flowers are developmentally similar with undifferentiated perianths composed of dense, tough tepals (high *FMA*; Figure 1). Many of these characteristics are common to other basal angiosperm lineages as well; trait variation for *Illicium* flowers (Figure 1, 3) is consistent with their having similarly costly flowers that maintain a hydraulic connection to the stem xylem (Feild et al. 2009). These common characteristics suggest that ANA grade and magnoliid flowers may rely predominantly on xylem delivery of water throughout anthesis to replace transpired water and maintain turgor. In order to maintain high transpiration rates and rely on xylem delivery of water throughout anthesis, basal angiosperm flowers have an efficient water transport system with highly branched veins and a high *K_flower_*. Significant trait and PIC correlations among the basal angiosperms suggest that these flowers may maintain water balance by coordinating water supply and loss over relatively short timescales, similar to angiosperm leaves (Figures 2,3).

In contrast, monocot and eudicot flowers exhibited little interspecific variation in hydraulic and structural traits and consistently had trait values associated with low fluxes of water and flowers with lower carbon investment. Although there were no significant relationships between stomatal traits and *K_flower_* (Table 2), stomatal abundance may nonetheless be important in determining the physiological strategies these flowers use. Stomata provide an open pathway for water inside the flower to evaporate into the atmosphere, and their near absence from many flowers may be an efficient way to prevent transpiration from tissues incapable of assimilating CO_2_. Stomatal traits were consistently and positively correlated with *g_min_*, suggesting that the near absence of stomata from monocot and eudicot flowers is critical in minimizing water loss. Furthermore, while angiosperm leaves can actively close their stomata to prevent desiccation (Brodribb & McAdam 2011; McAdam & Brodribb 2012), floral stomata may be limited in their ability to close (Hew et al. 1980) and, even when they do close, may not significantly curtail water loss (Teixido & Valladares 2014; Feild et al. 2009). Indeed, even under conditions that would induce stomatal closure, *g_min_* was lower among the monocots and eudicots than among basal angiosperm flowers (Figure 1, 2a). Among the monocots and eudicots, stomata may not play an important role in actively regulating water loss from flowers. Rather, water loss through the cuticle may have more strongly influenced the evolution of traits responsible for supplying water and maintaining turgor.

Reduced transpiration rates resulting from low stomatal abundance and low *g_min_* would have cascading consequences on other hydraulic and structural traits. Without the need to supply high fluxes of water to meet transpirational demands, constraints on the floral vascular system would be relaxed, allowing flowers to have a less ramified, less carbon intensive hydraulic system than those used by leaves, which have much longer functional lifespans (Roddy et al. 2013). Lower rates of water supply and loss would lengthen the turnover time of flower water, meaning that stored water (i.e. hydraulic capacitance) may be critical for maintaining turgor (Chapotin et al. 2003) and delaying desiccation (Figure 2b). Stored water may buffer water supply rates from water loss rates, which has been shown to occur in leafy shoots (Hunt & Nobel 1987) and which may explain why environmental conditions can have little impact on sap flow rates to flowers and inflorescences (Higuchi & Sakuratani 2005; Roddy & Dawson 2012). Because rates of water supply can be low, the phloem contribution of water might be a relatively larger fraction of the total water needed by the flower. In reality, flowers probably rely on a combination of water imported via the xylem and phloem and on stored water to maintain turgor throughout anthesis. The relative contributions of these sources, however, may be highly variable among species and throughout floral development.

## Conclusions

Flowers are one of the key innovations of the angiosperms and are incredibly diverse morphologically, yet the physiological costs of flowers can limit the extent to which floral morphology can be molded by pollinator selection. Flowers of the diverse monocot and eudicot clades have traits consistent with reduced rates of both water supply and water loss compared to basal angiosperms. This suggests that there may be substantial variation among these major clades in how flowers maintain turgor and prevent desiccation. One end of this continuum may be defined by maintaining a high hydraulic conductance to continuously supply water via the xylem, while the other end of the continuum may be defined by the reduction of water loss rates to delay desiccation. Reduced hydraulic conductance and greater reliance on stored water may physiologically buffer flowers from diurnal variability in the water status of other plant structures. Better understanding the mechanisms and timing of water transport to flowers and the tradeoffs between these mechanisms will be an important advancement in our understanding of floral physiology and its interaction with pollinator selection over evolutionary timescales. The present study establishes a comparative framework for characterizing the hydraulic mechanisms and strategies used by flowers.

## Acknowledgments

We thank R. Meinzer and two anonymous reviewers for useful comments on a previous draft of this manuscript, K.A. Simonin for insightful discussions, A. Schürkmann for technical assistance, and H. Forbes for facilitating access to plant material at the U.C. Botanic Garden. This work was funded by grants from the Jepson Herbarium and the Department of Integrative Biology, UC-Berkeley. A.B.R. was supported by a US NSF Graduate Fellowship and a postdoctoral fellowship from the Yale Institute for Biospheric Studies.

---

**Supplementary Figure S1.**
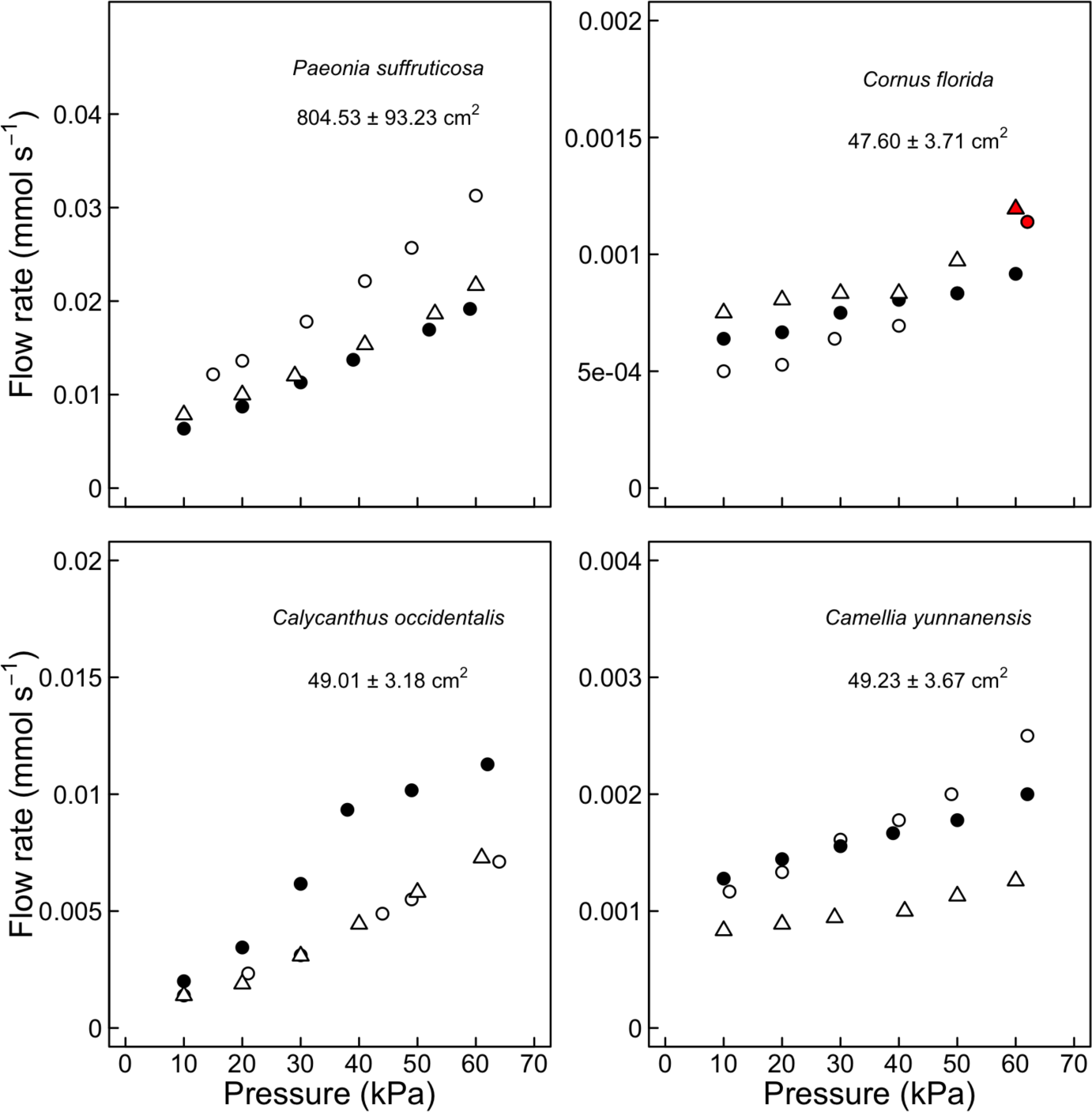

## References

Berendse F., Scheffer M. (2009) The angiosperm radiation revisited, an ecological explanation for Darwin’s ‘abominable mystery’. Ecology Letters 12, 865–872.

Brodribb T.J., Feild T.S. (2000) Stem hydraulic supply is linked to leaf photosynthetic capacity: Evidence from New Caledonian and Tasmanian rainforests. Plant, Cell & Environment 23, 1381–1388.

Brodribb T.J., Feild T.S., Jordan G.J. (2007) Leaf maximum photosynthetic rate and venation are linked by hydraulics. Plant Physiology 144, 1890–1898.

Brodribb T.J., Feild T.S., Sack L. (2010) Viewing leaf structure and evolution from a hydraulic perspective. Functional Plant Biology 37, 488.

Brodribb T.J., McAdam S.A. (2011) Passive origins of stomatal control in vascular plants. Science 331, 582–585.

Buckley T.N. (2015) The contributions of apoplastic, symplastic and gas phase pathways for water transport outside the bundle sheath in leaves. Plant, Cell & Environment 38, 7–22.

Chapotin S.M., Holbrook N.M., Morse S.R., Gutiérrez M.V. (2003) Water relations of tropical dry forest flowers: Pathways for water entry and the role of extracellular polysaccharides. Plant, Cell & Environment 26, 623–630.

Choat B., Gambetta G.A., Shackel K.A., Matthews M.A. (2009) Vascular function in grape berries across development and its relevance to apparent hydraulic isolation. Plant Physiology 151, 1677–1687.

Clearwater M.J., Luo Z., Ong S.E., Blattmann P., Thorp T.G. (2012) Vascular functioning and the water balance of ripening kiwifruit (*Actinidia chinensis*) berries. Journal of Experimental Botany 63, 1835–1847.

Clearwater M.J., Ong S.E.C., Li K.T. (2013) Sap flow and vascular functioning during fruit development. Acta Horticulturae 991, 385–392.

Costanza R., d’Arge R., De Groot R., Farber S., Grasso M., Hannon B., Limburg K., Naeem S., O’Neill R.V., et al. (1997) The value of the world’s ecosystem services and natural capital. Nature 387, 253–260.

Crepet W.L., Niklas K.J. (2009) Darwin’s second ‘abominable mystery’: Why are there so many angiosperm species? American Journal of Botany 96, 366–381.

Darwin C. (1888) The various contrivances by which orchids are fertilised by insects. John Murray, London.

Dodd M.E., Silvertown J., Chase M.W. (1999) Phylogenetic analysis of trait evolution and species diversity variation among angiosperm families. Evolution 53, 732–744.

Drake P.L., Price C.A., Poot P., Veneklaas E.J. (2015) Isometric partitioning of hydraulic conductance between leaves and stems: Balancing safety and efficiency in different growth forms and habitats. Plant, Cell & Environment 38, 1628–1636.

Feild T.S., Brodribb T.J. (2013) Hydraulic tuning of vein cell microstructure in the evolution of angiosperm venation networks. New phytologist 199, 720–726.

Feild T.S., Chatelet D.S., Brodribb T.J. (2009a) Ancestral xerophobia: A hypothesis on the whole plant ecophysiology of early angiosperms. Geobiology 7, 237–264.

Feild T.S., Chatelet D.S., Brodribb T.J. (2009b) Giant flowers of southern magnolia are hydrated by the xylem. Plant Physiology 150, 1587–1597.

Felsenstein J. (1985) Phylogenies and the comparative method. American Naturalist 125, 1–15.

Fenster C.B., Armbruster W.S., Wilson P., Dudash M.R., Thomson J.D. (2004) Pollination syndromes and floral specialization. Annual Review of Ecology, Evolution, and Systematics 35, 375–403.

Galen C. (1999) Why do flowers vary? Bioscience 49, 631–640.

Galen C., Dawson T.E., Stanton M.L. (1993) Carpels as leaves: Meeting the carbon cost of reproduction in an alpine buttercup. Oecologia 95, 187–193.

Galen C., Sherry R.A., Carroll A.B. (1999) Are flowers physiological sinks or faucets? Costs and correlates of water use by flowers of *Polemonium viscosum*. Oecologia 118, 461–470.

Grafen A. (1989) The phylogenetic regression. Philosophical Transactions of the Royal Society of London B 326, 119–157.

Greenspan M.D., Shackel K.A., Matthews M.A. (1994) Developmental changes in the diurnal water budget of the grape berry exposed to water deficits. Plant, Cell & Environment 17, 811–820.

Hetherington A.M., Woodward F.I. (2003) The role of stomata in sensing and driving environmental change. Nature 424, 901–908.

Hew C.S., Lee G.L., Wong S.C. (1980) Occurrence of non-functional stomata in the flowers of tropical orchids. Annals of Botany 46, 195–201.

Higuchi H., Sakuratani T. (2005) The sap flow in the peduncle of the mango (*Mangifera indica L.*) inflorescence as measured by the stem heat balance method. Journal of the Japanese Society of Horticultural Science 74, 109–114.

Higuchi H., Sakuratani T. (2006) Water dynamics in mango (*Mangifera indica L.*) fruit during the young and mature fruit seasons as measured by the stem heat balance method. Journal of the Japanese Society for Horticultural Science 75, 11–19.

Ho L.C., Grange R.I., Picken A.J. (1987) An analysis of the accumulation of water and dry matter in tomato fruit. Plant, Cell & Environment 10, 157–162.

Hunt E.R., Nobel P.S. (1987) Non-steady-state water flow for three desert perennials with different capacitances. Functional Plant Biology 14, 363–375.

John G.P., Scoffoni C., Sack L. (2013) Allometry of cells and tissues within leaves. American Journal of Botany 100, 1936–1948.

Kembel S.W., Cowan P.D., Helmus M.R., Cornwell W.K., Morlon H., Ackerly D.D., Blomberg S.P., Webb C.O. (2010) Picante: R tools for integrating phylogenies and ecology. Bioinformatics 26, 1463–1464.

Kerstiens G. (1996) Cuticular water permeability and its physiological significance. Journal of Experimental Botany 47, 1813–1832.

Kolb K.J., Sperry J.S., Lamont B.B. (1996) A method for measuring xylem hydraulic conductance and embolism in entire root and shoot systems. Journal of Experimental Botany 47, 1805–1810.

Kölreuter J.G. (1761) Vorläufige Nachrichten von einigen das Geschlecht der Pflanzen betreffenden Versuchen und Beobachtungen. Gleditschischen Handlung, Leipzig.

Lambrecht S.C. (2013) Floral water costs and size variation in the highly selfing *Leptosiphon bicolor* (Polemoniaceae). International Journal of Plant Sciences 174, 74–84.

Lang A. (1990) Xylem, phloem and transpiration flows in developing apple fruits. Journal of Experimental Botany 41, 645–651.

Lipayeva L.I. (1989) On the anatomy of petals in angiosperms. Botanicheskii Zhurnal 74, 9–18.

Li S., Zhang Y.J., Sack L., Scoffoni C., Ishida A., Chen Y.J., Cao K.F. (2013) The heterogeneity and spatial patterning of structure and physiology across the leaf surface in giant leaves of *Alocasia macrorrhiza*. PloS one 8, e66016.

McAdam S.A.M., Brodribb T.J. (2012) Stomatal innovation and the rise of seed plants. Ecology Letters 15, 1–8.

Memmott J., Waser N.M. (2002) Integration of alien plants into a native flower-pollinator visitation web. Proceedings of the Royal Society of London B. 269, 2395–2399.

Münch E. (1930) Die Stoffbewegungen in der Pflanze. Gustav Fischer, Jena, Germany.

Nobel P.S. (1983) Biophysical Plant Physiology and Ecology. W.H. Freeman, San Francisco.

Noblin X., Mahadevan L., Coomaraswamy I.A., Weitz D.A., Holbrook N.M., Zwieniecki M.A. (2008) Optimal vein density in artificial and real leaves. Proceedings of the National Academy of Sciences 105, 9140–9144.

Rasband W.S. (2012) <http://imagej.nih.gov/ij/> Bethesda, MD, USA: National Institutes of Health.

R Core Team. (2012) <http://www.R-project.org/> Vienna, Austria: R Foundation for Statistical Computing.

Roddy A.B., Dawson T.E. (2012) Determining the water dynamics of flowering using miniature sap flow sensors. Acta Horticulturae 951, 47–53.

Roddy A.B., Guilliams C.M., Lilittham T., Farmer J., Wormser V., Pham T., Fine P.V.A., Feild T.S., Dawson T.E. (2013) Uncorrelated evolution of leaf and petal venation patterns across the angiosperm phylogeny. Journal of Experimental Botany 64, 4081–4088.

Sack L., Cowan P.D., Jaikumar N., Holbrook N.M. (2003) The hydrology of leaves: Co¬ordination of structure and function in temperate woody species. Plant, Cell & Environment 26, 1343–1356.

Sack L., Dietrich E.M., Streeter C.M., Sánchez-Gómez D., Holbrook N.M. (2008) Leaf palmate venation and vascular redundancy confer tolerance of hydraulic disruption. Proceedings of the National Academy of Sciences 105, 1567–1572.

Sack L., Holbrook N.M. (2006) Leaf hydraulics. Annual Review of Plant Biology 57, 361–381.

Sack L., Scoffoni C., McKown A.D., Frole K., Rawls M., Havran J.C., Tran H., Tran T. (2012) Developmentally based scaling of leaf venation architecture explains global ecological patterns. Nature Communications 3, 837.

Savage J.A., Clearwater M.J., Haines D.F., Klein T., Mencuccini M., Sevanto S., Turgeon R., Zhang C. (*In press*.) Allocation, stress tolerance and carbon transport in plants: How does phloem physiology affect plant ecology? Plant, Cell & Environment.

Simonin K.A., Limm E.B., Dawson T.E. (2012) Hydraulic conductance of leaves correlates with leaf lifespan: Implications for lifetime carbon gain. New Phytologist 193, 939–947.

Skelton R.P., West A.G., Dawson T.E. (2015) Predicting plant vulnerability to drought in biodiverse regions using functional traits. Proceedings of the National Academy of Sciences 112, 5744–5749.

Sprengel C.K. (1793) Das entdeckte Geheimnis der Natur im Bau und in der Befruchtung der Blumen. Friedrich Vieweg dem aeltern, Berlin.

Stebbins G.L. (1970) Adaptive radiation of reproductive characteristics in angiosperms, I: Pollination mechanisms. Annual Review of Ecology and Systematics 307–326.

Strauss S.Y., Whittall J.B. (2006) Non-pollinator agents of selection on floral traits. In: Ecology and Evolution of Flowers. pp. 120–138. Oxford: Oxford University Press.

Teixido A.L., Valladares F. (2014) Disproportionate carbon and water maintenance costs of large corollas in hot Mediterranean ecosystems. Perspectives in Plant Ecology, Evolution and Systematics 16, 83–92.

Trolinder N.L., McMichael B.L., Upchurch D.R. (1993) Water relations of cotton flower petals and fruit. Plant, Cell & Environment 16, 755–760.

Webb C.O., Donoghue M.J. (2005) Phylomatic: Tree assembly for applied phylogenetics. Molecular Ecology Notes 5, 181–183.

Windt C.W., Gerkema E., Van As H. (2009) Most water in the tomato truss is imported through the xylem, not the phloem: A nuclear magnetic resonance flow imaging study. Plant Physiology 151, 830–842.

Zwieniecki M.A., Boyce C.K. (2014) Evolution of a unique anatomical precision in angiosperm leaf venation lifts constraints on vascular plant ecology. Proceedings of the Royal Society of London B. 281, 20132829.

